# Standardized brain and plasma EV enrichment pipeline validated for Single sample multi-Omic and fatty acids applications in Mouse and Human

**DOI:** 10.64898/2026.01.22.700328

**Authors:** Liam Barry-Carroll, Marjorie Varilh, Flore Marchaland, Chuck T Chen, Anne-Lisa Sadeyen, Jean-William Dupuy, Karina McDade, Tracey Millar, Richard P Bazinet, Sophie Layé, Anne-Aurelie Raymond, Alexandre Favereaux, Charlotte Madore, Jean-Christophe Delpech

**Affiliations:** University of Bordeaux, INRAE, Bordeaux INP, NutriNeuro, UMR 1286, F-33000 Bordeaux, France; Department of Nutritional Sciences, Faculty of Medicine, University of Toronto, Toronto, ON, Canada; Univ. Bordeaux, CNRS, INSERM, TBM-Core, US5, UAR 3427, OncoProt, F-33000 Bordeaux, France; University of Bordeaux, INSERM UMR1312, BoRdeaux Institute of onCology (BRIC), 33000, Bordeaux, France; Interdisciplinary Institute for Neuroscience, Centre National de la Recherche Scientifique, Bordeaux, France; Institute for Neuroscience and Cardiovascular Research, University of Edinburgh, Edinburgh, Scotland; Univ. Bordeaux, Bordeaux Proteome, F-33000 Bordeaux, France

## Abstract

Extracellular vesicles (EVs) are key mediators of intercellular communication, yet their molecular profiles across tissues and species remain poorly characterized, particularly due to currently available methods requiring a large amount of biological material (tissue or biofluids). Here, we established a workflow allowing the deep phenotyping of EV cargos starting from single samples of human and mouse origin. We took advantage of standardised EV isolation procedures and multi-omic techniques for the isolation and analysis of EVs from brain and plasma of human and mouse, integrating flow cytometric profiling, proteomics, miRNA sequencing, and fatty acid profiling. Here we report specific brain-derived EVs proteome, enriched in neuronal and glial proteins, polyunsaturated fatty acids profiles, and distinct miRNAs. At the periphery, we also report plasma-derived EVs signatures reflecting immune, metabolic, and systemic transport functions. Despite these expected material-specific differences, EVs from the same source displayed greater similarity across species than EVs from different material, supporting the translational relevance of mouse models. Importantly, using state-of-the-art miRNA profiling approach, we identified novel EV-specific miRNAs in human and mouse brain EVs, potentially allowing the exploration of new roles in neuronal signalling. Overall, we report here a method enabling deep multi-omic characterization from minimal starting material, offering a practical approach for studies with limited biological samples. These findings also demonstrate that the origin strongly shapes EV composition, highlighting conserved and species-specific molecular features, and provide a scalable framework for multi-omic investigations of EV biology.

**Summary Statement:** We present a standardised workflow allowing multi-omic profiling of brain and plasma-derived EVs from minimal human and mouse material. Our findings reveal both tissue-specific and species specific EV molecular signatures.

## Introduction

The interest and study of EVs has rapidly increased within most biological fields over the last years (Buzas, 2022; Cheng and Hill, 2022). However, this has coincided with numerous challenges, particularly in the development of methods for the successful and optimal isolation of EVs from various source material. A major challenge relates to the procurement of 100% pure EVs which is not always achievable due to overlaps in size and density of EVs with other particles and co-isolates (Brennan *et al*., 2020; Tian *et al*., 2020). Often the methods used for higher purity are also associated with several caveats. For example, the use of immunoaffinity capture methods to purify EVs based on the presence of specific surface proteins is increasingly popular, however this can ignore certain EV subsets due to the heterogeneous and unclear nature of any true ‘EV-markers’ (Tauro *et al*., 2012; Clayton *et al*., 2021). This is particularly important to consider when assessing different cell types. Indeed, another potential caveat of ultra-pure EV isolation is the low sample abundance which can affect and limit the downstream analyses of EVs using multi-omic techniques (Chitoiu *et al*., 2020; Shaba *et al*., 2022). A possibility to overcome this is to develop and use ultrasensitive methods, however these can be expensive and may not be readily available. Another option can include the pooling of several biological samples which is not always optimal for data interpretation (Webber and Clayton, 2013; Robinson *et al*., 2024).

Therefore, standardised methods and reporting are required which fit this balance between purity and abundance, to allow deep profiling of EV samples from individualized samples to keep sampling heterogeneity and biological relevance.

An often-unique feature of EV research is that a large proportion of the data has been recorded directly from human-derived biofluids in recent years (Chen *et al*., 2024). This is primarily owing to their ease of access and attractiveness as biomarkers in clinical studies (Conde-Vancells *et al*., 2010; Chen *et al*., 2024). However, this has led to a current gap within the field with regards to understanding the basic mechanisms of EV biology, as clinical and biomarker studies have outpaced fundamental studies from rodents. Evidently, there has been an increase in the use of *in vitro* studies to address such questions, particularly with the advent of readily available human induced pluripotent stem cell (iPSC) models (You *et al*., 2022). However, these do not always fully recapitulate the complexity of a living organism. Therefore, it is predicted that the utilisation of rodent models will be necessary to address this gap, especially as more sophisticated rodent models are developed. For this reason and for translation of basic research to clinical studies, it is important to understand whether the current methods used for EV isolation in humans can be effectively applied to rodent studies, to what degree there is an overlap in EV profile between both species and to evaluate the source specificity of EV signatures. Indeed, there are few studies which have compared EVs from mouse and human. Species-specific differences have been reported with regard to the miRNA content of plasma-derived EVs between humans and mice (Zhao *et al*., 2020). Additionally, comparative analysis of the proteome of brain-derived EVs from mouse and human have revealed so far only limited overlap in their EV signatures (Gallart-Palau, Serra and Sze, 2016). In regards to tissue or cellular source specificity of EVs, the use of plasma as a source of biomarkers for central nervous system (CNS) health has been the topic of multiple studies due to their predicted ability to pass between the blood brain barrier (Henriksen *et al*., 2007; Baird, Westwood and Lovestone, 2015; Banack *et al*., 2022; Nogueras-Ortiz *et al*., 2024). Furthermore the similarity between CNS and plasma-derived EVs should be addressed to estimate compatibility for use as source of blood-based biomarkers.

Here we customised commonly used and standardised methods for the isolation of EVs from tissue and fluid sources, and report an optimized EV analysis pipeline for multi-phenotyping from one individualized sample. In particular, we isolated EVs from brain tissue and plasma in both humans and mice. We successfully established an EV-dedicated workflow allowing the characterisation of EVs by transmission electron microscopy (TEM), flow cytometry, proteomic analysis, miRNA sequencing and profiling of polyunsaturated fatty acids starting from one human or mouse blood and brain sample. These analyses revealed EV-specific signatures related to protein and miRNA expression. A source-specific overlap was observed from EVs across mouse and human highlighting a degree of species conservation. Our analysis showed that while EVs from brain and plasma diverged, a number of proteins and miRNAs were also commonly shared. These data pinpoint both pan and source-specific profiles of EVs which will be of value in the study of EVs going forward.

## Results

### Isolation and identification of EV-enriched fractions from brain and plasma in human and mouse

Here we aimed to develop a simplified approach to isolate EVs from plasma and brain tissues of both human and mouse samples allowing a multitargeted omics analysis from the same specimen. Concerning the isolation of brain-derived EVs (BDEVs), non-fixed frozen mouse brains and blocks of human brain tissue of approximately 400 mg were used. Brain tissues were subjected to a combination of gentle mechanical homogenisation and enzymatic digestion to obtain a suspension of brain-derived cells. BDEVs were isolated from this cell suspension through a differential centrifugation approach followed by a sucrose-based density gradient in order to purify EVs based on their density as previously described (Vella *et al*., 2017; Muraoka *et al*., 2020) (**Figure 1 A**). Compared to the EV isolation procedure from biofluids, the methodology for extraction of EVs from tissue lacks standardisation. Therefore, we aimed to understand how the choice of digestive enzyme can alter the resultant EV extract. We compared the performance of papain and collagenase IV, two well-known enzymes, commonly used for the digestion of biological tissues, for BDEV isolation. For this comparison, whole mouse brains were subjected to the aforementioned BDEV isolation protocol, but using either papain or collagenase for the enzymatic digestion step. Seven fractions were collected (III-IX) from sucrose gradient. Each fraction was pelleted using ultracentrifugation and resuspended in filtered PBS. Nanoflow cytometric analysis of the resultant fractions revealed several differences in the properties of purified particles based on enzyme choice. Papain resulted in a significantly higher concentration of particles within fractions IV-VI (**Supplementary Figure 1 A**). Interestingly, there was also a decrease in the size of later fractions (VII and VIII) of papain treated samples and an increase in the free protein content present in papain treated samples compared to those isolated with collagenase (**Supplementary Figure 1 B, C**). Moreover, papain reduced the percentage of particles expressing the tetraspanin markers CD81, CD9 and CD63 compared to collagenase treated brains (**Supplementary Figure 1 D-F**). Altogether these findings indicate that collagenase IV is favourable for increasing the purity of preserved EVs isolated from mouse brain tissue. As such, collagenase IV was chosen for subsequent BDEV isolation from human and mouse brain tissue. Fractions V-VII were pooled and referred to as BDEVs due to their high concentration and expression of tetraspanins in both mouse and human, and subjected to downstream analyses (**Figure 1 B&C; Supplementary Figure 1 G-H**).

**Figure 1.**
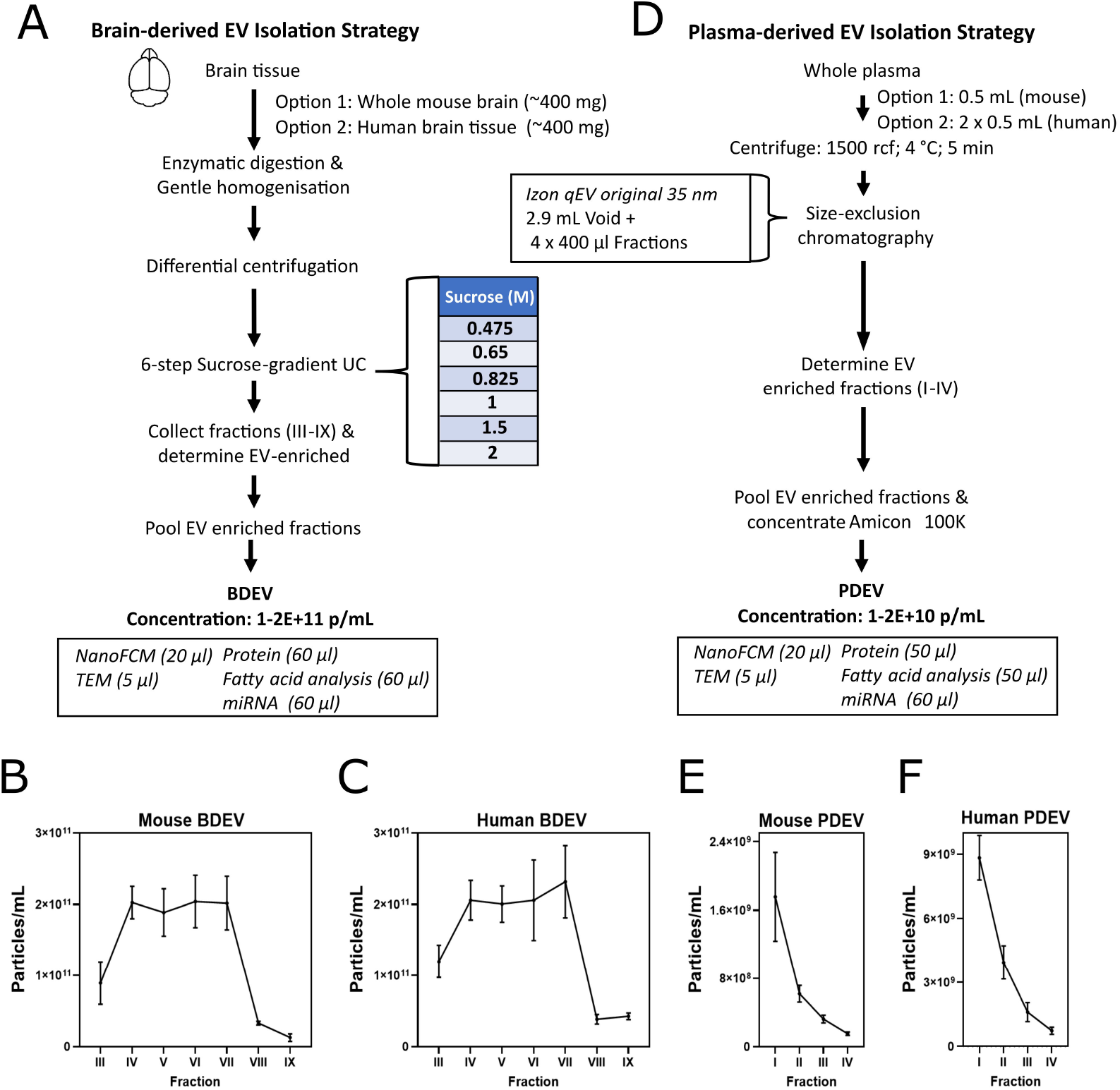
Isolation strategy of brain and plasma-derived extracellular vesicles. **(A)** Workflow of brain-derived EV isolation from whole mouse brain tissue and human brain punches utilising enzymatic digestion with homogenisation, differential centrifugation and a 6-steps sucrose-based density gradient. **(B-C)** Distribution of particle concentration across collected fractions from brain-derived EVs from mouse and human. **(D)** Workflow of a size-exclusion chromatography-based isolation of plasma-derived EVs from whole plasma. **(E-F)** Distribution of particle concentration across collected fractions from plasma-derived EVs from mouse and human. Data are expressed as mean ± SEM, n=4 per fraction.

For the isolation of plasma-derived EVs (PDEVs), both mouse and human plasma were subjected to a standard protocol of size-exclusion chromatography (SEC) (Brennan *et al*., 2020; Ter-Ovanesyan *et al*., 2021; Veerman *et al*., 2021). Owing to sample availability, 0.5 mL of plasma was used from a single mouse while 1 mL of plasma was used from human samples for EV analysis. 0.5 mL of plasma was applied to Izon’s qEV original 35 nm gen 2 columns. Four 400 µl fractions (I-IV) were eluted in filtered PBS following a void volume of 2.9 mL (**Figure 1 D**). Human plasma was applied to the column twice in 0.5 mL batches and the resulting fractions were pooled individually for the same sample. The identification of highly enriched EV fractions was assessed by measuring the concentration of particles and of free protein present in each fraction as commonly established (Gámez-Valero *et al*., 2016; Stranska *et al*., 2018; Ter-Ovanesyan *et al*., 2021). Indeed, for both mouse and human, fractions I and II contained the highest concentration of particles and lowest concentration of free protein resembling an EV enriched profile in accordance with previous studies (**Figure 1 E&F, Supplementary Figure 1 I-J**). Therefore, I and II fractions were pooled and subsequently concentrated using an 100Kda Amicon centrifugal filter to generate PDEVs and aliquoted for downstream analysis.

### Characterisation of BDEVs and PDEVs

The presence of particles displaying a typical EV ‘cup-like’ morphology could be observed using TEM negative staining of BDEV and PDEV samples from mouse and human (**Figure 2 A**). Analysis of the size distribution of BDEV particles up to 200 nm was conducted using the fiji plugin TEM ExosomeAnalyzer on acquired TEM images (Kotrbová *et al*., 2019) which revealed that mouse BDEVs had an average size of 134 nm, while human BDEVs had an average size of 127 nm (**Figure 2 B,C**). Further analysis of concentration and size was conducted using Nanoflow cytometry for particles ranging between 40-200 nm (**Figure 2 C-E**). The concentration of BDEVs and PDEVs was comparable between mouse and human (E+11 particles/ mL), however there was significant source-specific difference that was an order of magnitude between BDEVs and PDEVs from human and mouse (**Figure 2 C, D**). Similarly, there was no difference in size of EVs between species **(Figure 2 E**), although we observed a significant source-related difference, with smaller sized PDEVs compared to larger BDEVs only in mice (**Figure 2 E**). MPDEVs and HPDEVs had an average size of 75 nm and 75 nm, while MBDEVs and HBDEVs had an average size of 87 nm and 81 nm respectively (**Figure 2 C,E)**.

**Figure 1.**
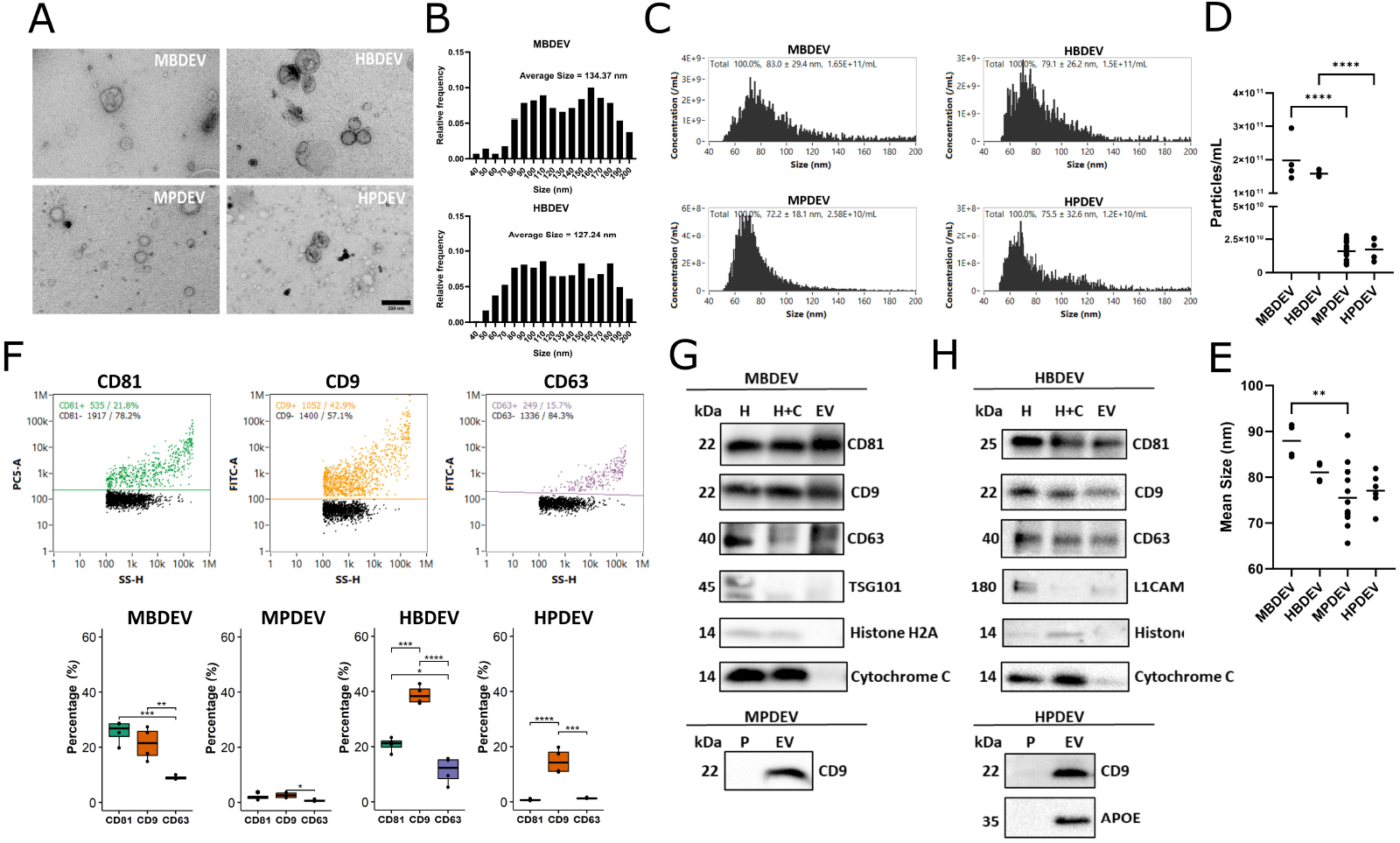
Characterisation of BDEV and PDEVs from human and mouse. **(A)** Representation of BDEV and PDEV from human and mouse by TEM. **(B)** Size distribution of BDEVs by TEM from human and mouse. **(C)** Representative size distribution of BDEVs and PDEVs from human and mouse by NanoFCM. **(D-E)** Comparison of mean concentration and size of BDEV and PDEVs (MBDEV=4, MPDEV=13, HBDEV=4, HPDEV=6). **(F)** Representative dot plots displaying CD81+, CD9+ and CD63+ events from HBDEVs. Analysis of CD81, CD9 and CD63 expression (%) in BDEVs and PDEVs by NanoFCM (n=4). **(G-H)** Representative immunoblots of commonly detected proteins in EVs (CD81, CD9, CD63, TSG101 and L1CAM) and negative control proteins (Histone H2A, Cytochrome C and APOE) in BDEV and PDEVs from human and mouse (H=Homogenate, H+C=Homogenate + Collagenase IV, P=Plasma) . Data are expressed as mean ± SEM, One-way ANOVA followed by Bonferroni’s post-hoc analysis for multiple (F) and pairwise comparisons (D and E). Statistical differences: *p<0.05, **p<0.01, ***p<0.001, ****p<0.0001. Scale bar is 200 nm.

Next, the tetraspanin phenotype of EVs was assessed by Nanoflow cytometry, and we observed that the signature of CD81, CD9 and CD63 was highly variable between EV source and species. BDEVs from mouse had a significantly higher expression of CD81 compared with CD9 and CD63, whereas HBDEVs contained a significantly higher expression of CD9 compared with CD81 and CD63 (**Figure 2 F**). A common observation was that CD63 was the least abundant tetraspanin in BDEVs in both mouse and human. CD9 was the most abundant tetraspanin found in PDEVs from mouse and human (**Figure 2 F**). The protein expression of CD81, CD9 and CD63 was observed in EVs isolated from brain of mouse and human. These markers were also found in brain homogenate fractions with and without collagenase from mouse and human (**Figure 2 G-H**), as expected as it contains free EVs and intracellular pools of EVs. Notably, the expression of non-EV markers Cytochrome C and Histone H2A were absent in EV fractions from both mouse and human brain. PDEVs were characterised by their expression of CD9 and the expression of APOE was also observed in PDEVs from human highlighting the presence of lipoprotein as co-isolates of PDEVs using SEC (**Figure 2 H**). Interestingly, CD81 and CD63 were not detected in PDEVs by western immunoblot (data not shown) in both human and mouse. Altogether our characterisation of EVs from brain and plasma supports EV identity and is in agreement with the MISEV guidelines (Welsh et al., 2024).

### Proteomic profiling of BDEVs and PDEVs reveal EV-specific signatures

In order to investigate the proteomic landscape of our EVs, high sensitivity mass-spectrometry was performed from just 1 ug of protein using the search algorithm CHIMERYS. To decode proteins that were specific to EVs from brain and plasma, we compared the relative abundance between the EV samples and respective reference controls (brain lysate for BDEVs and plasma for PDEVs) (**Table S1**). For whole mouse brain, 41 proteins were significantly enriched in mouse BDEVs (MBDEVs) whereas 167 proteins were found to be enriched in mouse brain lysate (**Figure 3 A; Table S2**). Top proteins significantly enriched in MBDEV included membrane-associated proteins, proteins involved in protein trafficking and sorting, and EV biogenesis, including CLDND1, MIB1, ARHGEF101, and ATP6AP2, all of which have been previously reported in EVs (Burke *et al*., 2014; Ikeda *et al*., 2021; Murray *et al*., 2021; Ye *et al*., 2025) (**Figure 3 A, B**). In contrast, the potassium channel subunit KCNJ16 may represent a novel marker of MBDEVs. Proteins that were enriched in mouse brain lysate included the ribonuclear protein POP1 (Michelson *et al*., 2024), a regulator of autophagosome formation PIK3C3 (Wei *et al*., 2025), the histone protein H1-4 and SNCA, which is involved in synaptic vesicle trafficking (Carnazza *et al*., 2022). From human brain, 24 proteins were found to be significantly enriched in human BDEVs (HBDEVs), while 30 proteins were found significantly enriched in the brain lysate (**Figure 3 C, Table S2**). Most significantly enriched proteins found in HBDEVs included KLHL42, UBL7, PITPNM2, and DHRS11, which are associated with membrane trafficking (Grabon, Bankaitis and McDermott, 2019), ubiquitin-like signalling (Lear *et al*., 2020; Jiang *et al*., 2023), and lipid metabolism (Endo *et al*., 2020) and may represent novel markers of HBDEVs (**Figure 3 C, D, Table S2**). Human brain lysate specific proteins included PRPH, AKR7L and SLC39A10, which are related to axonal growth (Bjornsdottir *et al*., 2019), oxidoreductase activity (Rajib and Sharif Siam, 2020) and zinc transport (Kawata *et al*., 2025).

**Figure 3.**
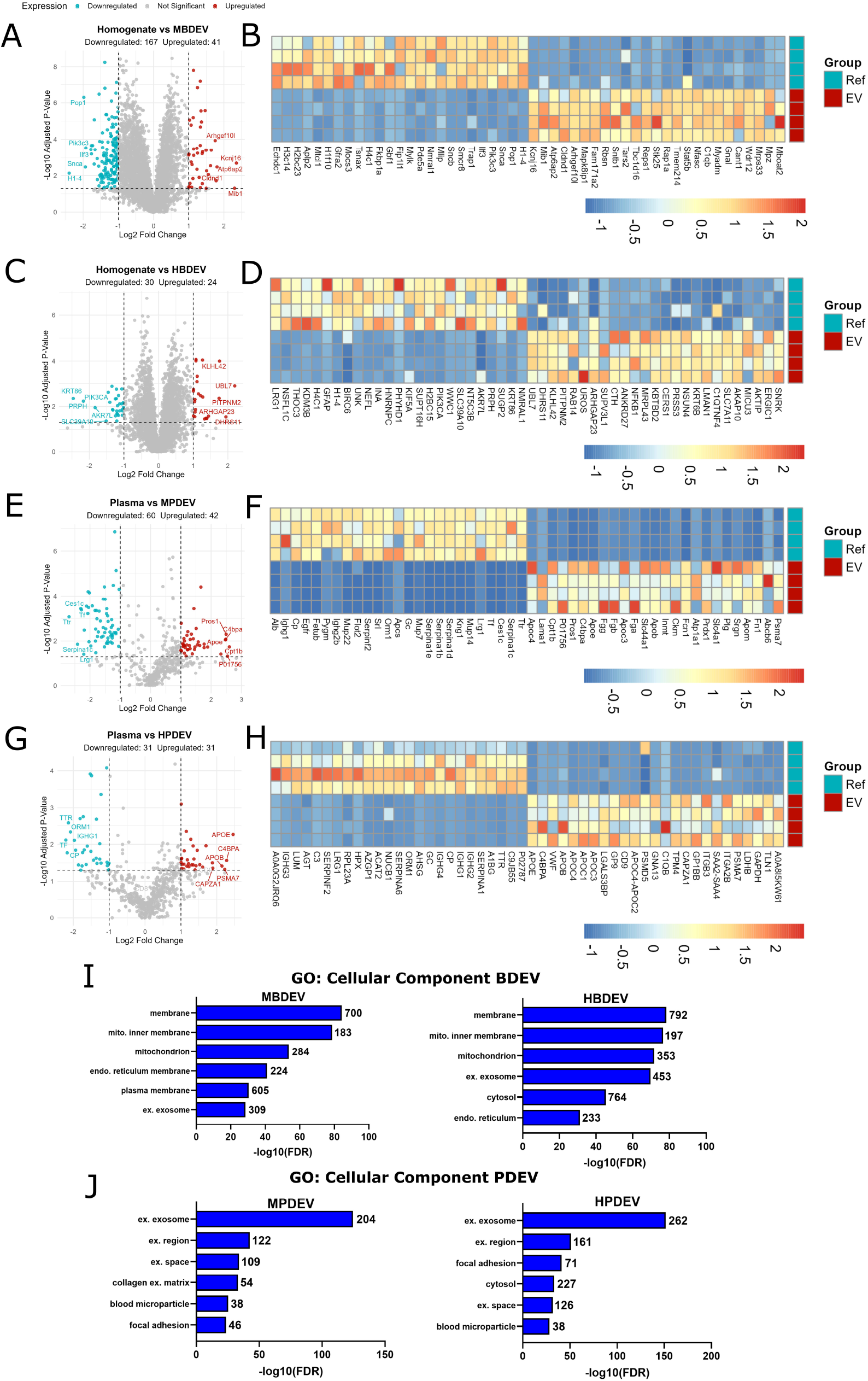
Identification of EV specific protein signature from brain and plasma. **(A)** Volcano plot comparing protein enrichment between BDEVs and the brain homogenate samples from mouse. For volcano plots, the log_2_ fold change (x-axis) represents the relative enrichment of proteins in extracellular vesicles compared to the reference sample, while the -log_10_(p-value) (y-axis) indicates statistical significance. Proteins with significant enrichment (adjusted p-value < 0.05) are highlighted in colour, with those enriched in extracellular vesicles shown in red and those enriched in the reference sample in blue. Non-significant enrichment proteins are shown in grey. Dotted lines represent thresholds for statistical significance and biological relevance. Key proteins of interest are labelled. **(B)** Heatmap of the top 25 enriched proteins found in BDEVs and brain homogenate samples from mouse. The heatmaps display the Z-score normalized abundance of the most enriched proteins in EVs and the reference sample. Rows represent individual proteins, while columns correspond to biological replicates. Colour intensity reflects relative protein abundance, with red indicating higher expression and blue indicating lower expression. **(C)** Volcano plots comparing protein enrichment between PDEVs and the plasma samples from mouse. **(D)** Heatmap of the top 25 enriched proteins found in PDEVs and plasma samples from mouse. **(E)** Volcano plots comparing protein enrichment between BDEVs and the brain homogenate samples from human. **(F)** Heatmap of the top 25 enriched proteins found in BDEVs and brain homogenate samples from human. **(G)** Volcano plots comparing protein enrichment between PDEVs and the plasma samples from human. **(H)** Heatmap of the top 25 enriched proteins found in PDEVs and plasma samples from human. **(I-J)** Top 6 GO: Cellular component terms associated with the EV protein signature (protein enrichment ratio >1.5) from mouse and human brain and mouse and human plasma. GO analysis was performed using DAVID and bar plot displays the top six enriched GO terms, ranked by statistical significance. The y-axis represents the GO terms, while the x-axis shows the -log_10_(FDR), where higher values indicate stronger enrichment.

From mouse plasma, 42 proteins were found to be significantly enriched in MPDEVs compared to plasma, while 60 were found significantly enriched in plasma (**Figure 3 E, Table S2**). Top significantly enriched proteins from MPDEVs were primarily proteins associated with membrane, immune function and lipid metabolism and have been previously reported in EVs (Moreira-Costa *et al*., 2021; Du *et al*., 2022; Jiang *et al*., 2024). These included PROS1, C4BPA and APOE, while CPT1B has not been explicitly reported in PDEVs (**Figure 3 F, F, Table S2**). The top hits from mouse plasma included many abundant soluble proteins such as CES1C, TF, TTR, and SERPINA1C (Kuiperij *et al*., 2009; Gan *et al*., 2023; Wan *et al*., 2025). From human plasma, 31 proteins were significantly enriched in HPDEVs and 31 were found specific to plasma (**Figure 3 G, Table S2**). While some of the top HPDEV proteins were also reported in MPDEVs, CAPZA1, C4BPA, APOE, APOB, and PSMA7 proteins were associated to cytoskeletal remodelling, lipid transport and proteostasis (Zheng *et al*., 2017; Wang *et al*., 2021; Du *et al*., 2022; Jiang *et al*., 2024) (**Figure 3 G, H, Table S2**). APOE and APOB may highlight the potential co-isolation of lipoproteins which are abundant in plasma, although they have also been reported to be EV-related (Jiang *et al*., 2024).

Next, we performed an analysis of gene ontology (GO) terms for cellular component using the human orthologues for identified mouse proteins, due to the lower level of annotation of EV-related proteins in murine samples (**Supplementary Figure 2 A**). Overall, the protein signature of both MBDEV and HBDEV was associated with the terms membrane and included extracellular exosome (**Figure 3 I**). Interestingly, canonical markers of EVs, including CD9, CD81, and CD63, were found to be enriched in BDEVs from both human and mouse samples (**Table S1**). Altogether these findings show that the proteome of BDEVs is enriched with functionally relevant signalling and membrane-associated proteins that can distinguish them from intracellular/autophagosome proteins. The top hit from GO: Cellular component analysis from the most abundant proteins for both MPDEV and HPDEV was extracellular exosome (**Figure 3 J, Table S2**). Similar to mouse, human plasma was enriched with abundant circulating proteins. Together these findings highlight an enrichment of EVs in our plasma-derived samples and suggest that there is a strong overlap in proteome between species. Indeed, we see that EVs derived from the same tissue type—whether brain or plasma—exhibit greater similarity across species than EVs from different tissues within the same organism with 67.1% of proteins shared between MBDEV and HBDEV, and 31.3% of proteins shared between MPDEV and HPDEV (**Supplementary Figure 2 C, D**). This is compared to just 9.8% of shared proteins between MBDEV and MPDEV and 17.7% of proteins shared between HBDEV and HPDEV (**Supplementary Figure 2 E, F**).

### BDEV and PDEV overlap with reported EV signatures and exhibit divergent cell-type enrichment

Next we aimed to compare the protein signature from our plasma and brain EVs with a well-characterised EV-specific database, Exocarta. For this, proteins with an enrichment ratio greater than 1.5 from mouse and human EVs were compared with the reported human Exocarta proteins list (**Table S3**). The human proteins from Exocarta were chosen for comparison due to the very low number of EV-specific proteins reported in mouse samples (**Supplementary Figure 2 A**). Overall there is a substantial overlap between Exocarta and our EV-enriched protein signature from both brain and plasma (**Figure 4 A-D**). However, a higher percentage of proteins from PDEVs overlapped with Exocarta compared with that of BDEVs (**Figure 4 A-D**). Cellular component pathway analysis with DAVID revealed GO: Extracellular exosome as the top hit for all overlapping proteins between Exocarta and EV-enriched proteins from brain and plasma (**Figure 4 E**). Conversely, the EV-enriched proteins not found in Exocarta were related to the GO: mitochondrial inner membrane for both mouse and human brain-derived EVs (**Figure 4 F**). Non-overlapping proteins from plasma-derived EVs were related to GO: Immunoglobulin complex in mouse and GO: High-density lipoprotein particle in human (**Figure 4 F**). Differences between the number of Exocarta-associated proteins from plasma and brain may stem from the reportedly higher number of studies reporting protein signature from PDEVs compared to BDEVs.

**Figure 4.**
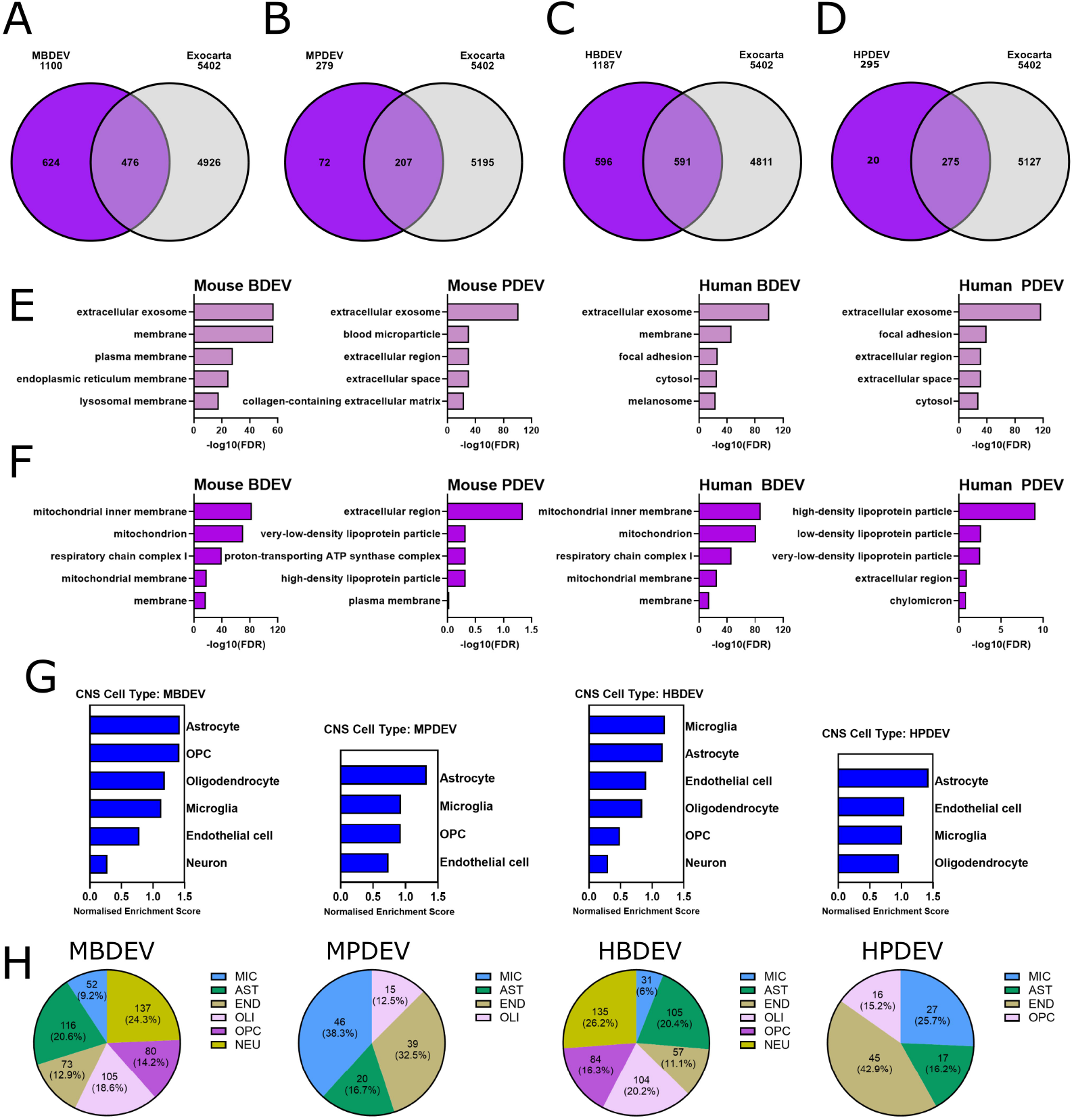
Unravelling the EV protein signature from brain and plasma. **(A-D)** Overlap of EV protein signature from mouse brain, mouse plasma, human brain and human plasma with the human Exocarta protein database. **(E)** Top 5 GO: cellular component terms associated with the shared proteins between Exocarta and EVs from mouse brain, mouse plasma, human brain and human plasma. **(F)** Top 5 GO: cellular component terms associated with the EV proteins that are not reported in Exocarta mouse brain, mouse plasma, human brain and human plasma. GO analysis was performed using DAVID. The bar plot displays the top five enriched GO terms, ranked by statistical significance. The y-axis represents the GO terms, while the x-axis shows the -log_10_(FDR), where higher values indicate stronger enrichment. The log_2_ fold change (x-axis) represents the relative enrichment of proteins in extracellular vesicles compared to the reference sample, while the -log_10_(p-value) (y-axis) indicates statistical significance. **(G)** GSEA CNS cell type enrichment analysis of MBDEV, MPDEV, HBDEV and HPDEV related proteins using the McKenzie et al., 2018 dataset to estimate potential cells of origin. The bar plot displays the overrepresented (blue) CNS cell types, ranked by normalised enrichment score. **(H)** Estimation of the proportion proteins from MBDEVs, MPDEVs, HBDEVs and HPDEVs related to microglia (MIC), astrocytes (AST), endothelial cells (END), oligodendrocytes (OLI), oligodendrocyte precursor cells (OPC) and neurons (NEU).

In order to estimate the cells of origin from our EVs, GSEA cell type enrichment analysis was conducted based on the EV-enriched protein signature for all of our samples and the human cell landscape and mouse cell atlas were selected as reference databases (Elizarraras *et al*., 2024) (**Supplementary Figure 2 G**). Immune cells were the most common between EV sources and species with neutrophils found in MBDEVs, T cells reported in MPDEVs, HBDEVs and HPDEVs. Myeloid cells were found to be enriched in HBDEVs and HPDEVs and monocytes were enriched in MPDEVs (**Supplementary Figure 2 G)**. As expected, cells of brain origin were enriched in both MBDEVs and HBDEVs including oligodendrocytes, astrocytes and foetal neurons (**Supplementary Figure 2 G**). Next, the EV-enriched protein signature from all sources were compared with the profile of the most abundant CNS cell types including microglia, astrocytes, oligodendrocytes, OPCs, endothelial cells and neurons (McKenzie *et al*., 2018). GSEA analysis was conducted in order to reveal the most enriched CNS cell types based on the protein signature from brain and plasma which considers the enrichment factor of proteins in our samples (**Figure 4 G**). This revealed that the protein signature from MBDEV and HBDEV were enriched for all cell types (**Figure 4 G-H**). Notably there were some species differences where MBDEVS top enriched CNS cell type was astrocyte, whereas HBDEVs were mostly enriched for microglia. Interestingly, neurons were the least enriched CNS cell type for both MBDEVs and HBDEVs. Plasma-derived EVs from both mouse and human were not enriched for neurons and there were some species differences (**Figure 4 G-H**). The protein signature of MPDEVs were enriched for astrocytes, microglia, OPC and endothelial cells, whereas HPDEVs were significantly enriched for astrocytes. OPCs were not enriched in HPDEVs whereas endothelial cells and microglia were (**Figure 4 G-H**).

### Assessing the miRNA content of BDEVs and PDEVs

We then analysed the miRNA content of EVs and reference sample from brain and plasma in both mice and humans in order to identify EV-specific hits using a newly available high sensitivity method (**Table S4**). For analysis of miRNA in EV samples, 60 µl of sample containing a range of 5.6E+9 - 4.91E+11 particles/mL were used. A significant correlation was observed between the number of miRNA reads and concentration of BDEVs, whereas this relationship was not found for PDEVs (**Supplementary Figure 3 A-B**), suggesting that the packaging of miRNA content is different between PDEVs and BDEVs. Out of 409 miRNAs detected, MBDEVs had 2 significantly upregulated miRNAs and 4 which were significantly downregulated and specific to the homogenate control (**Figure 5 A-B, Table S2**). The 2 MBDEV enriched miRNAs were mmu-miR-6239 which has not been previously described in brain EVs and may present novel BDEV markers, and mmu-miR-6240 which has been previously detected in murine mast cells (Hu *et al*., 2021). MPDEVs showed a more distinct signature with 22 upregulated and 27 downregulated miRNAs compared with plasma (**Figure 5 C-D, Table S2**), out of 493 miRNAs detected. MPDEV-enriched miRNAs included let-7e-5p, mmu-miR-409-5p, mmu-miR-153-3p, mmu-miR-541-3p, and mmu-miR-467d-5p, which have been reported in rodent EVs and may play a role in inflammatory processes (Hu *et al*., 2021; Cao *et al*., 2022; Xue *et al*., 2024). Overall HBDEVs showed the most pronounced EV-enriched miRNA profile with 157 upregulated and 32 downregulated miRNAs out of 618 (**Figure 5 E-F, Table S2**), of which miR-331-3p, miR-193-3p, miR-129-2-3p, miR-324-3p, or miR-1908-5p were among the top hits of HBDEV-specific miRNAs. Interestingly, none of these have been previously reported in EVs which again may highlight novel EV-specific miRNAs. Finally, out of 375, 38 miRNAs were upregulated in HPDEVs, while 10 were downregulated (**Figure 5 G-H, Table S2**). Our observation that miR-129-5p, miR-195-5p, miR-125b-5p, miR-138-5p, and likely let-7c-5p, are enriched in HPDEVs is supported by existing literature which shows EV packaging of these miRNAs (Raimondo *et al*., 2020; Xun *et al*., 2021; Yang *et al*., 2021). While direct evidence for miR-138-5p and let-7c-5p in plasma EVs is limited, their detection in human EVs and prevalence of the let-7 family in plasma strongly support their relevance. Out of the detected miRNAs from both MBDEVs and MPDEVs, 21% were common between both EV types although PCA analysis showed a source specific distinction (**Supplementary Figure 3 C-D**). A similar source-specific profile and overlap was observed in human samples with 22% of detected miRNAs shared between HBDEVs and HPDEVs (**Supplementary Figure 3 E-F**). Previous studies have sought to define sequence features associated with miRNA enrichment in EVs, leading to the identification of short RNA motifs termed EXOmotifs (Villarroya-Beltri *et al*., 2013; Santangelo *et al*., 2016; Garcia-Martin *et al*., 2022). Analysis of previously described EXOmotifs within the miRNAs enriched in HBDEV revealed that 82 of the 193 HBDEV-enriched miRNAs contained at least one known EXOmotif (**Supplementary Figure 4 A**). Because these motifs were originally characterized in non-neuronal cell types, we next performed an unbiased motif discovery analysis to identify brain-specific EXOmotifs associated with HBDEV. Using the MEME Suite (https://meme-suite.org/meme/), we identified a putative novel brain EXOmotif, CCA/UG, which was present in 40.86% of HBDEV-enriched miRNAs (**Supplementary Figure 4 B**). This motif may represent a previously unrecognized sequence signature contributing to miRNA sorting into brain-derived EVs. Altogether, these findings suggest species-specific differences in the enrichment of EV miRNAs, particularly when comparing BDEVs.

**Figure 2.**
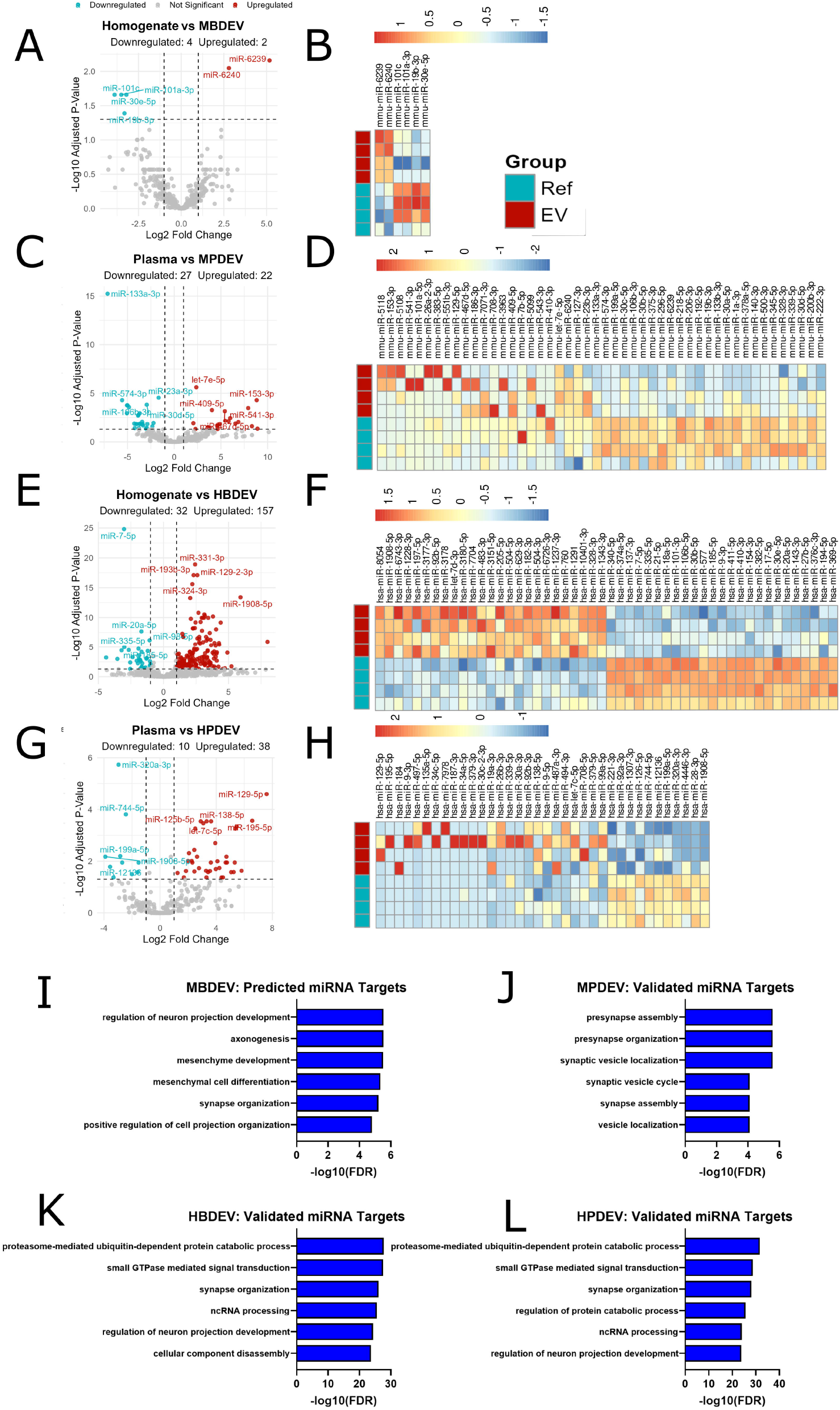
Upregulated miRNAs in EVs from brain and plasma. **(A)** Volcano plot of both downregulated and upregulated miRNAs from BDEVs and brain homogenate from mouse. Downregulated miRNAs were considered with >-1 Log2 Fold change and >1.25 of -Log10 of adjusted P value. Upregulated miRNAs were considered with >1 Log2 Fold change and >1.25 of -Log10 of adjusted P value. The log_2_ fold change (x-axis) represents the relative expression of miRNAs in EVs compared to the reference sample, while the -log_10_(p-value) (y-axis) indicates statistical significance. miRNAs with significant enrichment (adjusted p-value < 0.05) are highlighted in colour, with those enriched in EVs shown in red and those enriched in the reference sample in blue. Non-significant enrichment proteins are shown in grey. Dotted lines represent thresholds for statistical significance and biological relevance. Upregulated miRNAs of interest are labelled. **(B)** Heatmap of the top upregulated and downregulated miRNAs in BDEVs and brain homogenate samples from mouse. The heatmaps display the Z-score of normalised expression. Rows represent individual proteins, while columns correspond to biological replicates. Colour intensity reflects relative protein abundance, with red indicating higher expression and blue indicating lower expression. **(C)** Volcano plots comparing miRNA expression between PDEVs and the plasma samples from mouse. **(D)** Heatmap of the top expressed miRNAs found in PDEVs and plasma samples from mouse. **(E)** Volcano plots comparing miRNA expression between BDEVs and the brain homogenate samples from human. **(F)** Heatmap of the top expressed miRNAs found in BDEVs and brain homogenate samples from human. **(G)** Volcano plots comparing miRNA expression between PDEVs and plasma samples from human. **(H)** Heatmap of the top expressed miRNAs in PDEVs and plasma samples from human. **(I-L)** Top 6 miRNA targets for MBDEVs, MPDEVs, HBDEVs and HPDEVs. Target analysis was performed using mulitMIR. Bar plot displays the top six targets, ranked by statistical significance. The y-axis represents the targets, while the x-axis shows the -log_10_(FDR), where higher values indicate stronger enrichment.

Next, in order to determine the potential functions of these miRNAs, we performed target analysis of upregulated miRNAs found across different EVs. Of note, there were no validated targets of MBDEV-miRNA, only predicted, which may highlight the novelty of these MBDEV-miRNAs. The biological pathways (BPs) related to the predicted targets of the MBDEV-miRNAs were primarily related to neuronal functioning, and included regulation of neuron projection development, axonegenesis and synapse organization (**Figure 5 I**). Validated targets of MPDEVs were screened and we found BPs related to synaptic function and vesicle trafficking (**Figure 5 J**). Interestingly there was a larger overlap in the top BPs related to the validated targets of miRNAs from HBDEVs and HPDEVs (**Figure 5 K, L**). Proteasome mediated ubiquitin-dependent protein catabolic process, small GTPase mediated signal transduction and synapse organization were the top 3 BPs for targets of both HBDEV and HPDEV miRNAs (**Figure 5 K, L**). The frequency of pathways related to synaptic function may highlight a conserved function of EV-miRNAs across compartment and species. Indeed, several studies have highlighted the potential function of miRNAs in synaptic plasticity (Cohen *et al*., 2011; Ye *et al*., 2016).

### Probing the fatty-acid profile of EVs

Fatty acids (FA) such as palmitic acid (PA), arachidonic acid (ARA), docosahexaenoic acid (DHA), docosapentaenoic acid (DPA) n-3 and n-6 are found esterified in the glycerol backbones of phospholipids, a main component of the lipid bilayer. Considering EVs are formed primarily of lipid membrane bilayer, we characterise these FA components. Overall, our targeted FA analysis revealed that BDEVs from mouse and human contained a higher concentration of the measured FAs compared with PDEVs (**Figure 6 A**). This may be due to source material, which is a whole brain and is known to be rich in membranous structures. Next, we assessed the proportion of these five targeted FAs from EVs, and found that BDEVs from both mouse and human share a very similar overlap in FA signature, primarily containing PA, ARA and DHA with a small amount of DPA n-3 and n-6 (**Figure 6 B**). Conversely, PDEVs had the highest percentage of PA followed by ARA in human and mouse (**Figure 6 B**). Interestingly MPDEVs also contained a portion of DHA that was not reflected in HPDEVs (**Figure 6 B**). We also found that concentration of detected FAs seem to be proportional to the starting EV concentration, as it was significant for certain FAs including ARA, DHA and DPA n-3 (**Figure 6 C-E**), but not for DPA n-6 or PA (**Figure 6 F,G**). This may relate to the more pronounced role of ARA, DHA and DPA n-3 in membrane integrity, whereas DPA n-6 and PA are reported to be less implicated.

**Figure 6.**
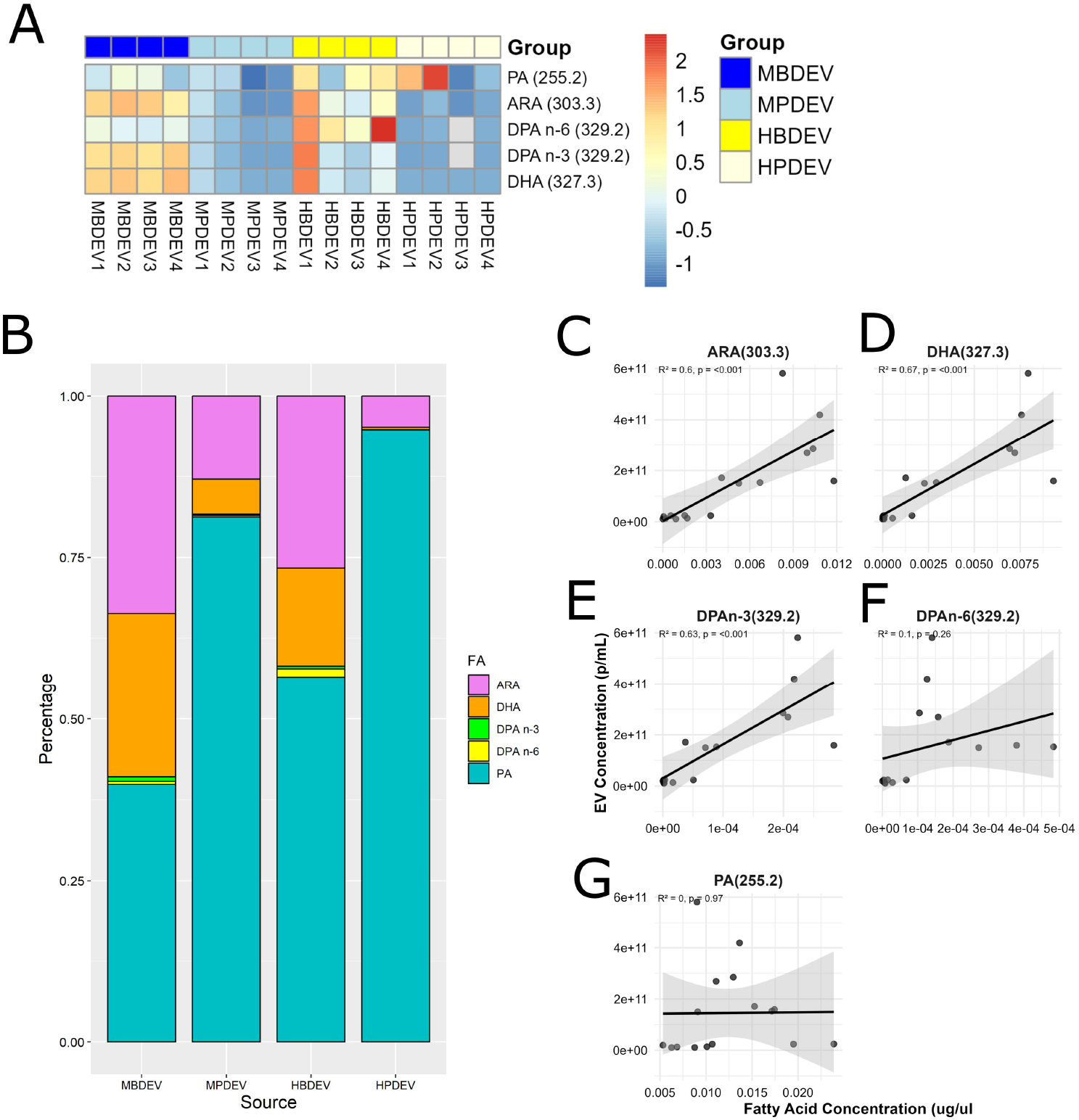
The FA profile of EVs from brain and plasma. **(A)** Heatmap of the z-score of fatty acid concentrations of palmitic acid (PA), arachidonic acid (ARA), docosapentaenoic acid (DPA) n-6, DPA n-3 and docosahexaenoic acid (DHA) from mouse and human BDEVs and PDEVs. **(B)** Cumulative plot of the overall fatty acid profile from MBDEVs, MPDEVs, HBDEVs and HPDEVs. **(C-G)** Linear regression of the relationship between the starting concentration of EVs and resultant concentration of the ARA, DHA, DPA n-3, DPA n-6 and PA.

## Discussion

In this study, we customised existing protocols for the isolation of EVs from plasma and brain tissue of mice and humans, while adhering to MISEV guidelines (Welsh et al., 2024). We performed a deep comparison of these EVs, employing a range of biochemical and molecular techniques to characterize their broad physical properties, proteome, miRNA cargo, and fatty acid composition as recommended by MISEV guidelines which demonstrated an enrichment of EVs in our samples. The findings here provide insight into how EV origin and species influence EV composition and underscore the importance of source material in shaping vesicle identity.

An interesting finding is that there is a source-specific heterogeneity of EVs between brain and plasma. This was consistently observed across the different analyses performed, including the biophysical properties of particles as well as the content of EVs revealed by proteomic profiling, miRNA sequencing, and fatty-acid analysis. It is likely that the method of EV isolation and the nature (liquid or tissue) of starting material also contributes to these differences. Overall, our findings suggest that the tissue of origin plays a dominant role in dictating EV molecular composition, while species-specific factors contribute comparatively to smaller variations. This cross-species similarity has been previously found in BDEVs and supports the utility of mouse models for studying EV biology, especially when focusing on brain-derived EVs. However, it also highlights the necessity of carefully matching tissue sources when translating findings from animal models to human systems. This source-specific EV signature likely reflects both the unique cellular composition and physiological functions of the brain and plasma environments. Indeed, BDEVs were enriched in neuronal and glial proteins, CNS-enriched miRNAs and fatty acids typical of neural membranes (e.g., DHA, ARA) (Salem Jr. *et al*., 2001; Sambra *et al*., 2021). In contrast, PDEVs showed higher representation of proteins linked to immune signalling, metabolic regulation, and systemic transport, as well as a distinct lipid profile more consistent with peripheral cell origin. While we applied standardized protocols across species and tissues, it should be noted that different isolation approaches used between BDEVs and PDEVs can enrich for different subtypes of vesicles or co-isolate contaminants, potentially influencing interpretation (Gámez-Valero *et al*., 2016).

Our study also addresses a practical challenge in EV biology: the limited availability of starting material from brain tissue and clinical biofluids (Newman, Useckaite and Rowland, 2022; Cross *et al*., 2024; Escudero-Cernuda *et al*., 2025). Multi-omic analyses were successfully performed on single EV preparations from mouse and human BDEVs, as well as from human PDEVs, while the combination of EV preparations from two animals were needed to complete the full analysis of mouse PDEVs. By explicitly detailing the input requirements for each workflow, we demonstrate a strategy to maximize information gain from limited material amount, an issue particularly relevant for translational and clinical studies. Our protocol enables efficient multi-omic characterization without compromising data quality, providing a framework for researchers working with scarce biological material. A critical advantage of our workflow is the ability to conduct deep multi-omic profiling from only 1 µg of EV protein, corresponding to ∼500 µL of plasma or ∼400 mg of brain tissue per sample. This yielded over 1,000 proteins from plasma EVs and more than 6,000 proteins from brain EVs from a single isolation. By comparison, many plasma EV proteomic studies require 1–2 mL of plasma and typically identify 300–800 proteins (Karimi *et al*., 2018; Reymond, Gruaz and Sanchez, 2023; Fochtman *et al*., 2024), with higher yields often dependent on pooling or additional depletion steps. Recent methodological advances, such as optimized data-independent acquisition (DIA) mass spectrometry, have further demonstrated the feasibility of deep proteome coverage with ultra-low input. Cross et al. (2024), for example, reported the detection of ∼3,800 proteins from as little as ∼50 ng of EV peptides (∼1 µg protein input), in line with the depth achieved in our study (Cross *et al*., 2024)

Our miRNA analyses further reinforce the tissue-driven specificity of EV molecular cargo while also uncovering potentially novel vesicle-associated small RNAs. BDEVs demonstrated strong correlations between vesicle concentration and miRNA read depth, suggesting a more uniform packaging of small RNAs in cerebral EVs compared to the heterogeneous profiles observed in PDEVs. This observation aligns with prior studies showing that neuronal activity and synaptic vesicle trafficking can directly influence the selective incorporation of miRNAs into EVs (Cohen *et al*., 2011; Ye *et al*., 2016). Notably, we identified several BDEV-enriched miRNAs not previously reported in the EV literature, including miR-6239, miR-6240, and a subset of human-specific candidates such as miR-331-3p and miR-129-2-3p. Their predicted or validated targets were strongly enriched in pathways related to synapse organization, axonogenesis, and vesicle trafficking, suggesting potential functional roles in neuronal circuit regulation. PDEVs, by contrast, showed signatures more consistent with immune and systemic signaling, in agreement with prior reports describing circulating EV miRNAs as markers of inflammation and metabolic homeostasis (Raimondo *et al*., 2020; Xun *et al*., 2021; Yang *et al*., 2021). Importantly, the overlap of top biological processes targeted by human BDEV and PDEV miRNAs—particularly proteasome-mediated protein catabolism, small GTPase signaling, and synapse organization—suggests that certain EV-mediated regulatory pathways may be conserved across compartments and species. Finally, cumulative demonstrations have been made on the role of EXOmotifs as sorting sequences that determine EV secretion from different cell types, even though the mechanisms by which miRNAs are sorted or retained in their cells of origin remains largely unknown. Most of the results have been obtained on peripheral cell types including white and brown adipocytes, endothelium, liver and muscle, with the beginning of the identification of cell-specific sorting sequences, without ruling out the positivity of shared motifs depending on the tissue of origin (Villarroya-Beltri *et al*., 2013; Santangelo *et al*., 2016; Garcia-Martin *et al*., 2022). Exploring EXOmotifs in our brain specific datasets, the observation of the presence of previously identified EXOmotifs in 43% of the HBDEVs and the newly identified brain-specific EXOmotif in 40% of the HBDEVs are very interesting and strengthen the need for future research on these particular motifs and their contribution to the secretion of BDEVs. This could ultimately participate in future research aiming to improve targeting in RNA-mediated therapies.

The fatty acid composition of EVs, even if limited in our study to 5 FAs, also reflected tissue-of-origin signatures. Both mouse and human BDEVs were enriched in DHA and ARA, fatty acids known to be highly concentrated in neuronal membranes where they support membrane fluidity, signaling, and synaptic function (Salem Jr. *et al*., 2001; Valenzuela *et al*., 2021). PDEVs, on the other hand, were dominated by palmitic acid (PA) with relatively lower contributions from DHA and ARA, consistent with their derivation from peripheral cells (Abdelmagid *et al*., 2015). Interestingly, we observed a species-specific divergence in plasma vesicles, with mouse PDEVs retaining detectable DHA that was absent in human PDEVs, potentially reflecting differences in lipid metabolism between rodents and humans. Correlation analyses further revealed that the abundance of DHA, ARA, and DPA n-3 scaled with EV concentration, whereas PA and DPA n-6 did not, supporting the view that these long-chain fatty acids are integral components of EV membranes rather than contaminants or loosely associated species. These results parallel earlier lipidomic characterizations of Evs that highlighted the enrichment of polyunsaturated fatty acids in brain- and non-brain-derived vesicles (Llorente *et al*., 2013; Skotland *et al*., 2017, 2020; Su *et al*., 2021).

Taken together, our results establish one of the most comprehensive multi-omic profiles of brain and plasma EVs to date from human and mouse species, while also providing a practical roadmap for overcoming the sample quantity barriers that often limit EV research. By integrating proteomic, miRNA, and lipidomic analyses across species and tissues, this work not only advances understanding of EV biology by revealing new EV specific markers, but also contributes a scalable framework for future studies aimed at translational biomarker discovery and mechanistic insight.

### Methods Animals

Male C57BL6/J mice (Janvier laboratories, Le Genest-Saint-Isle, France) were kept on a 12-h light/12-h dark schedule, housed on poplar woods chips litter in a controlled environment (21-23°C, 40% humidity), with access to water and chow ad libitum. All animal procedures followed the guidelines of the EU Directive 2010/63/EU for animal experiments and received approval from the national ethical committee for animal care and use (approval ID 39219).

### Human tissues and plasma

All post-mortem tissues were collected *via* the Edinburgh Brain Bank (ethics approval from East of Scotland Research Ethics Service, 21/ES/0087) in line with the Human Tissue (Scotland) Act 2006. The use of human tissue for post-mortem studies has been reviewed and approved by the Edinburgh Brain Bank ethics committee and approved by institutional review board of Inserm (IRB/ CEEI – avis #20-748). Samples included posterior hippocampus (n=3) and prefrontal cortex (n=1) from male subjects aged 69-84. Human blood samples were extracted from each donor at the EFS, Service Ressources biologiques, Bordeaux, France by venipuncture and collected into EDTA coated tubes (∼5 ml).

### Isolation of brain-derived EVs

Whole frozen mouse brain or human cortical brain tissue pieces (approximately 400 mg) were dissected into thin strips of around 0.3 cm x 1 cm on ice. For enzymatic digestion, the dissected tissue was incubated with 3.2 mL of prewarmed (37°C) Hibernate E solution containing either 20 units of papain (Worthington LK003176) or 200 units of collagenase IV (Worthington). The samples were incubated at 37°C for a total of 20 minutes including gentle shaking at 5 minutes and trituration with a 25 mL pipette tip at 15 minutes. At 20 minutes, the enzymatic reaction was stopped by placing the samples on ice and addition of 6 mL of Hibernate E containing protease/phosphatase inhibitor (1X) (Fisher). The resultant homogenate was gently dissociated into a single cell suspension using a glass potter and subsequently passed through a 40 µm cell strainer. The filtrate was subjected to differential centrifugation at 4°C: 300 x g (10 min), 2000 x g (10 min) and 100,000 x g (70 min) with retention of the pellet at each step and a final filtration step of the supernatant using a 40 µm strainer before centrifugation at 100,000 x g. The resultant pellet containing EVs was resuspended in 2 mL of 0.475 M of sucrose and applied to the top of a sucrose gradient containing 5 layers of sucrose at increasing molarity: 0.65 M, 0.825 M, 1 M, 1.5 M and 2 M (top to bottom). The sucrose gradient was centrifuged for 20 h at 200,000 x g; 4°C (SW-41-Ti swing bucket rotor, Beckman Coulter). Gradient fractions were recovered as follows: III (1 mL), IV (2 mL), V (2 mL), VI (2 mL), VII (2 mL), VIII (2 mL) and IX (1 mL). Each fraction was diluted with filtered PBS up to 10 mL and centrifuged (100,000 x g; 4°C; 7 min). Pellets were resuspended in 40 µl of filtered PBS.

### Isolation of plasma-derived EVs

Mouse blood was collected by intracardiac puncture and transferred into an EDTA coated Eppendorf tube (∼1.5 ml). Plasma was obtained by centrifuging whole blood at room temperature for 15 min at 1,600g. The plasma samples were stored at −80 °C. EVs were isolated from the plasma of either 3 month old male mice, or male donors aged 60-70 yrs old. For both species, frozen plasma was thawed and centrifuged at 1500 xg for 5 mins at 4 °C to remove any debris. After being brought to room temperature, 500 µl of plasma was applied to an equilibrated IZON original 35nm size-exclusion chromatography column. After a void volume of 2.9 mL, 4 fractions of 500 µl were eluted in filtered PBS and collected for further analysis. This process was repeated to twice with a total of 1 mL of plasma. Fractions I-II were subsequently pooled from both runs and concentrated using an Amicon 100K 2mL filter device. The final volume was recorded and samples were aliquoted for downstream analyses.

### Western Immunoblotting

For assessment of the protein content, samples were lysed on ice with TENT buffer (50 mM Tris HCl [pH 7.5], 2 mM EDTA, 150 mM NaCl, 1% Triton X-100) for 15 mins and sonicated (3 x 1 sec pulses). Lysates were then clarified by centrifugation (3600 rpm; 5 min; 4°C) and the protein concentration in each sample will be determined using a BCA assay (Interchim). Lysates were equalized in TENT buffer and denatured in 2X Lamelli buffer (BioRad) at 95 °C for 5 mins without the addition of a reducing agent. A total of 15 µg of MBDEVs and HBDEVs protein was loaded or 15 µl of MPDEV and HPDEV protein lysate was loaded on to 4–20% TGX Stain Free gels (BioRad) and then transferred onto nitrocellulose membranes (BioRad). Membranes were blocked at RT for 1 h with blocking solution (5% non-fat milk diluted in TBST) and then probed with primary antibodies (1:1000) diluted in blocking solution overnight at 4°C. The following primary antibodies were used: CD9 (Merck, CBL162); CD63 (BioOrbyt, orb229829); TSG101 (Santa Cruz Biotechnology, sc-7964); β-Actin (BioLegend, 622102); Cytochrome C (Cell Signaling Technology/Ozyme, 11940); Histone H2A.Z (Cell Signaling Technology/Ozyme, #2718); CD81 (Cell Signaling Technology/Ozyme, 10037); CD81 (Cell Signaling Technology/Ozyme, 56039); Alix (Cell Signaling Technology/Ozyme, #2717S); CD9 (Cell Signaling Technology/Ozyme, 98327); CD63 (Ozyme, CAB5271); L1CAM (Abcam, ab24345); APOE (Cell Signaling Technology, #13366). The membranes were further incubated with horseradish peroxidase conjugated secondary antibody (1:5000) for 1 h. To allow for visualization of total protein and ensure equal loading in the absence of an exosomal housekeeping protein, gels (Biorad) were activated by UV light to allow trihalo compounds within the gel to react with tryptophan residues of proteins in a UV-induced reaction to produce fluorescence which was visualized using a BioRad ChemiDoc™ MP imaging system (stain-free method). Immunoreactivity was detected using SuperSignal™ West Dura enhanced chemiluminescence (ECL) solution (ThermoFisher).

### Negative Staining & Transmission Electron Microscopy

An aliquot of EV sample was fixed with 4% paraformaldehyde on ice. The fixed samples (10 μl) were absorbed onto a glow-discharged 300-mesh heavy duty carbon-coated formvar Cu grids (ProSciTech, Kirwan, QLD) for 20 min followed by washing with PBS (3 x 5 min). The grids were then incubated with 1% glutaraldehyde for 5 min and washed with MilliQ water (8 x 2 min). Subsequent negative staining was carried out by incubating the grids with uranyl-oxalate in the dark for 5 mins followed by incubation with methyl cellulose-uranyl acid solution in the dark and on ice for 10 mins. High-contrast images were acquired using a HITACHI H7650 transmission electron microscope (Hitachi, Krefeld, Germany). The negative staining and electron microscopy was performed at the Bordeaux Imaging Centre.

### NanoFCM

For analysis of size, concentration and antigenic phenotype of EVs, a Flow NanoAnalyzer (NanoFCM) containing 488⍰nm and a 638⍰nm laser lines was calibrated using 200⍰nm polystyrene beads (NanoFCM) with a defined concentration of 2.168⍰×⍰1010 particles⍰/ml, which also served as a reference for particle concentration. In addition, a cocktail of silica beads (NanoFCM) of four different diameters (68⍰nm, 9⍰1nm, ⍰3⍰nm and 155⍰nm) served as size reference standards. EV samples were diluted with filtered PBS allowing for the detection of between 2000 and 14000 events in 1 minute. Particle concentration and size distribution were calculated using NanoFCM software (NF Profession V2.0). For staining, 4 µl of undiluted sample was incubated with 1 µl of fluorescently conjugated antibodies (APC anti-CD9 (Biolegend;), APC anti-CD63 (Biolegend;), PE anti-CD81 (Biolegend;), APC anti-CD9 (Biolegend;), APC anti-CD63 (Biolegend;), APC anti-CD81 (Biolegend;)) for 30 minutes at room temperature. The fluorescent membrane dye MemGlow488 (Universal Biologicals;) was prediluted (1:100) with filtered PBS and 4 µl of undiluted sample was incubated with 1 µl of diluted MemGlow.

### Mass spectrometry-based label-free quantitative proteomics

Four independent biological replicates for each experimental condition were used. *in-gel* protein digestion by the trypsin was performed as previously described (Henriet *et al*., 2017). NanoLC-MS/MS analysis were performed using an Ultimate 3000 RSLC Nano-UPHLC system (Thermo Scientific, USA) coupled to a nanospray Orbitrap Fusion™ Lumos™ Tribrid™ Mass Spectrometer (Thermo Fisher Scientific, California, USA). Each peptide extracts were loaded on a 300 µm ID x 5 mm PepMap C_18_ precolumn (Thermo Scientific, USA) at a flow rate of 10 µL/min. After a 3 min desalting step, peptides were separated on a 50 cm EasySpray column (75 µm ID, 2 µm C_18_ beads, 100 Å pore size, ES903, Thermo Fisher Scientific) with a 4-40% linear gradient of solvent B (0.1% formic acid in 80% ACN) in 57 min. The separation flow rate was set at 300 nL/min. The mass spectrometer operated in positive ion mode at a 2.0 kV needle voltage. Data was acquired using Xcalibur 4.4 software in a data-dependent mode. MS scans (m/z 375-1500) were recorded at a resolution of R = 120000 (@m/z 200), a standard AGC target and an injection time in automatic mode, followed by a top speed duty cycle of up to 3 seconds for MS/MS acquisition. Precursor ions (2 to 7 charge states) were isolated in the quadrupole with a mass window of 1.6 Th and fragmented with HCD@28% normalized collision energy. MS/MS data was acquired in the Orbitrap cell with a resolution of R=30000 (@m/z 200), a standard AGC target and a maximum injection time in automatic mode. Selected precursors were excluded for 60 seconds. Protein identification and Label-Free Quantification (LFQ) were done in Proteome Discoverer 3.1. CHIMERYS node using prediction model inferys_3.0.0 fragmentation was used for protein identification in batch mode by searching against a Uniprot *Homo sapiens* database (82415 entries, release January, 2024) or a UniProt *Mus musculus* database (55052 entries, release January, 2024). Two missed enzyme cleavages were allowed for the trypsin. Peptide lengths of 7-30 amino acids, a maximum of 3 modifications, charges between 2-4 and 20 ppm for fragment mass tolerance were set. Oxidation (M) and carbamidomethyl (C) were respectively searched as dynamic and static modifications by the CHIMERYS software. Only “high confidence” peptides were retained corresponding to a 1% false discovery rate at peptide level. Minora feature detector node (LFQ) was used along with the feature mapper and precursor ions quantifier. The normalization parameters were selected as follows: (1) Unique peptides, (2) Precursor abundance based on intensity, (3) Normalization mode: total peptide amount, (4) Protein abundance calculation: summed abundances, (5) Protein ratio calculation: pairwise ratio based and (6) Missing values are replaced with random values sampled from the lower 5% of detected values. Quantitative data were considered for master proteins, quantified by a minimum of 2 unique peptides, a fold changes ≥ 2 and a statistical p-value adjusted using *Benjamini-Hochberg* correction for the FDR lower than 0.05. The mass spectrometry proteomics data have been deposited to the ProteomeXchange Consortium via the PRIDE (Deutsch *et al*., 2023) partner repository with the dataset identifiers: PXD069631 (Human) and PXD069905 (Mouse).

In the case of BDEVs, brain lysate was used as the reference control, whereas for PDEVs, plasma was used. Following protein identification and relative quantification of abundance, the protein signatures of EVs and reference samples were revealed based on statistical significance and protein enrichment ratio which was calculated on the protein abundance in EV compared to the reference sample.

### Fatty acid profiling of EVs from brain and plasma

Total lipids were extracted from samples according to the method of Folch, Lees, and Sloane-Stanley (Folch et al., 1957) using a mixture of chloroform:methanol (2:1 by volume). The mixture was centrifuged at 1470g for 10 mins at room temperature and the chloroform layer were then extracted. New chloroform was added to the remaining aqueous phase and the mixture was once again centrifuged. The chloroform layer was extracted and added to the previously extracted chloroform layer. This total lipid extract (TLE) was evaporated under nitrogen and reconstituted in a known volume of chloroform. A known amount of heptadecanoic acid (17:0), used as internal standard, was added to the TLE. TLE was then saponified with 10% potassium hydroxide in methanol. Saponified total fatty acids were converted to pentafluorobenzyl esters by heating at 60°C in 1:10:1,000 (v:v:v) pentafluorobenzyl bromide:diisopropylamine:acetonitrile, dried under nitrogen gas, and dissolved in heptane for GC-MS analysis. They were analysed on an Agilent 5977A quadrupole mass spectrometer coupled to an Agilent 7890B gas chromatograph (Agilent Technologies, Mississauga, ON, Canada) in negative chemical ionization mode. Peaks were identified on the basis of selected fragmented ions.

### miRNA analysis and sequencing

Total RNA was extracted from extracellular vesicle (EV) preparations using the miRNeasy Serum/Plasma Kit (Qiagen), following the manufacturer’s protocol. RNA quality and concentration were assessed using a NanoDrop spectrophotometer and an Agilent 2100 Bioanalyzer. Small RNA libraries were constructed using the NEXTflex Small RNA-Seq Kit v3 (PerkinElmer), as previously described (Soula et al., 2018). Briefly, 1–5 µg of total RNA was ligated with 3’ and 5’ adapters, followed by reverse transcription and amplification to generate cDNA libraries. The libraries were size-selected to enrich for small RNA species, including microRNAs. Sequencing was performed on an Illumina platform, and data were analysed to identify and quantify miRNAs.

### Statistical analysis

All statistical analyses were performed using the latest version of GraphPad Prism (GraphPad Software). Data are presented as mean ± SEM unless otherwise stated, and statistical significance was assessed using the appropriate parametric or non-parametric tests, with p<0.05 considered significant. Gene ontology (GO) Cellular Component enrichment analyses of proteomic data were performed using the Database for Annotation, Visualization and Integrated Discovery (DAVID). Results were visualised based on -log10(FDR). Where FDR was equal to 0, half of the lowest non-zero FDR value in the same dataset was inputted for visual representation. Gene Set Enrichment Analysis (GSEA) on complete mouse and human cell atlases was carried out using the online WEB-based GEne SeT AnaLysis Toolkit (Elizarraras *et al*., 2024), while GSEA analysis based on CNS cell types was carried out using the most recent release of the GSEA application (Broad Institute) using a brain cell type specific gene dataset of astrocytes, endothelial cells, microglia, neurons, oligodendrocytes and OPCs (McKenzie et al., 2018) against the enriched EV proteins using the protein enrichment ratio for ranking. Predicted and validated miRNA targets were identified using the multiMIR R package, and subsequent functional enrichment of miRNA targets was analysed with Bioconductor packages for GO analysis.

## Acknowledgements

We thank the Bordeaux Imaging Center for the TEM microscopy, a service unit of the CNRS-INSERM and Bordeaux University, member of the national infrastructure France BioImaging supported by the French National Research Agency (ANR-10-INBS-04) with the help and guidance from Melina Petrel and Sabrina Lacomme. We also thank the EFS, Service Ressources biologiques, Bordeaux, France, for their help in human blood donor incorporation in the manuscript. The authors also wish to thank staff from the Centre d’IngénieRie ComportementalE on Bordeaux’s Neurocampus (CIRCE) and the GPR Brain for its support of the CIRCE. Finally, we would like to thank the RRI Food4BrainHealth network, in which SL, RPB, CC, CM and JCD are members, for the scientific opportunities it provides.

## Funding

This work was funded in part by the Région Nouvelle-Aquitaine CHESS Exomarquage 13059720-13062120 (JCD, SL), the ANR-23-CE14-0056-01 (JCD), the Postagreenskills fund INRAE PAF14_Omine (CM), the human nutrition INRAE department young researcher starting grant (JCD, CM).

## Author contributions

Conceptualization, J.C.D., CM.; Methodology, J.C.D., C.M., L.B.C., A.A.R., A.F., C.T.C., R.B.; Investigation, L.B.C., M.V., F.M., C.T.C., A.L.S., J.W.D.; Writing – Original Draft, J.C.D.,C.M., L.B.C.; Writing – Review & Editing, all authors; Formal Analysis: J.C.D., C.M., L.B.C., C.T.C., F.M., J.W.D., A.F.; Funding Acquisition, J.C.D., C.M., S.L., R.B., A.F.; Supervision, J.C.D., C.M.

## Author information

To be added upon request

## Conflict of Interest

No conflict of interest to declare.

## Figure Legends

**Supplementary Figure 1.**
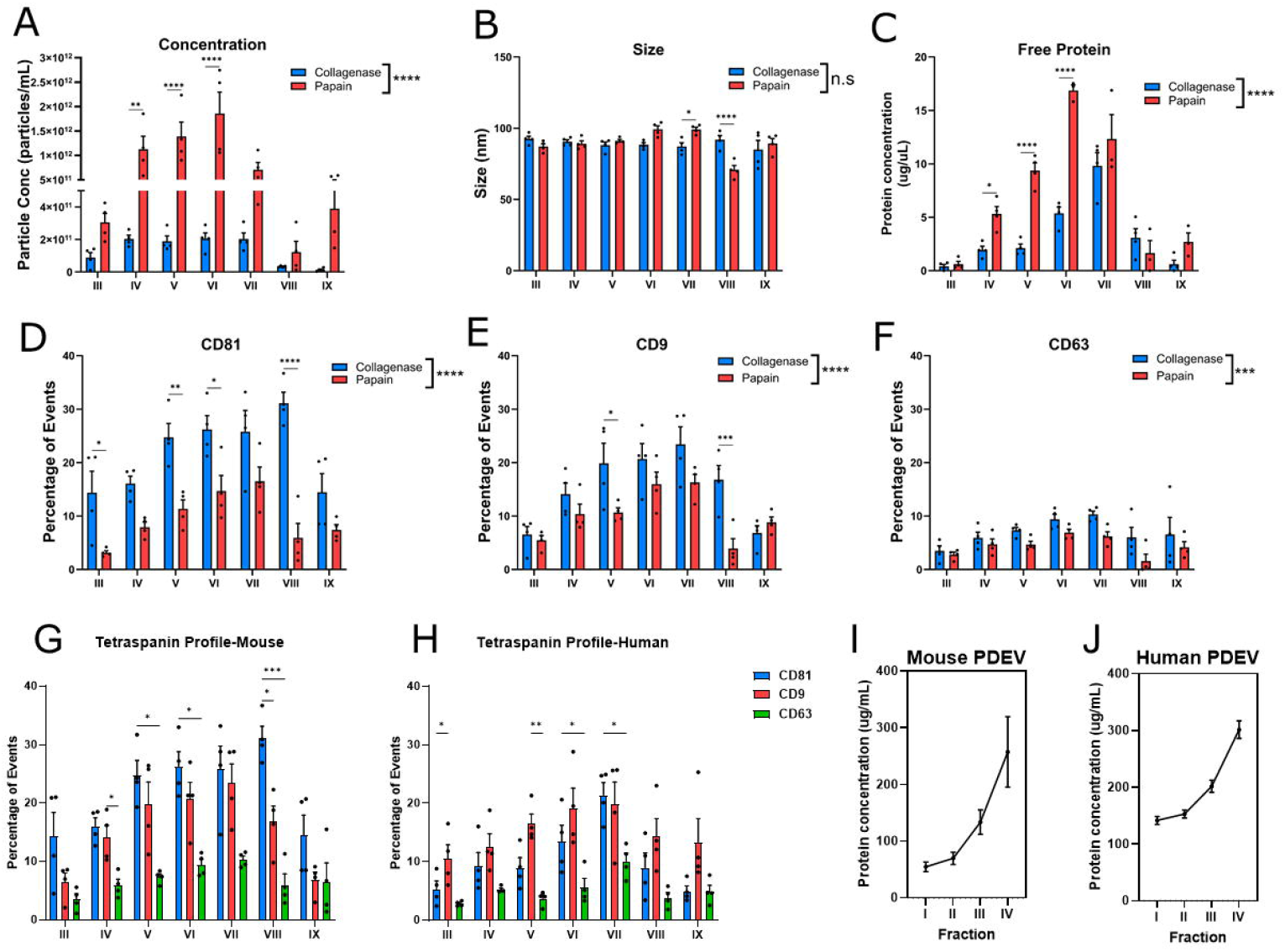
**(A-C)** Comparison of average particle concentration, size and free protein concentration from fractions III-IX between collagenase and papain treated mouse brains. **(D-F)** Comparison of the percentage of CD81^+^, CD9^+^ and CD63^+^ events in fractions III-IX between collagenase and papain treated mouse brains. **(G-H)** Comparison between the percentage of CD81^+^, CD9^+^ and CD63^+^ events in fractions III-IX from mouse and human **(I-J)** Free protein concentration (ug/mL) in fractions I-IV of plasma EV isolates in mouse and human. Data are expressed as mean ± SEM, collagenase n=4, papain n=4, Two-way ANOVA followed by Bonferroni’s post-hoc analysis. Statistical differences: *p<0.05, **p<0.01, ***p<0.001, ****p<0.0001.

**Supplementary Figure 2.**
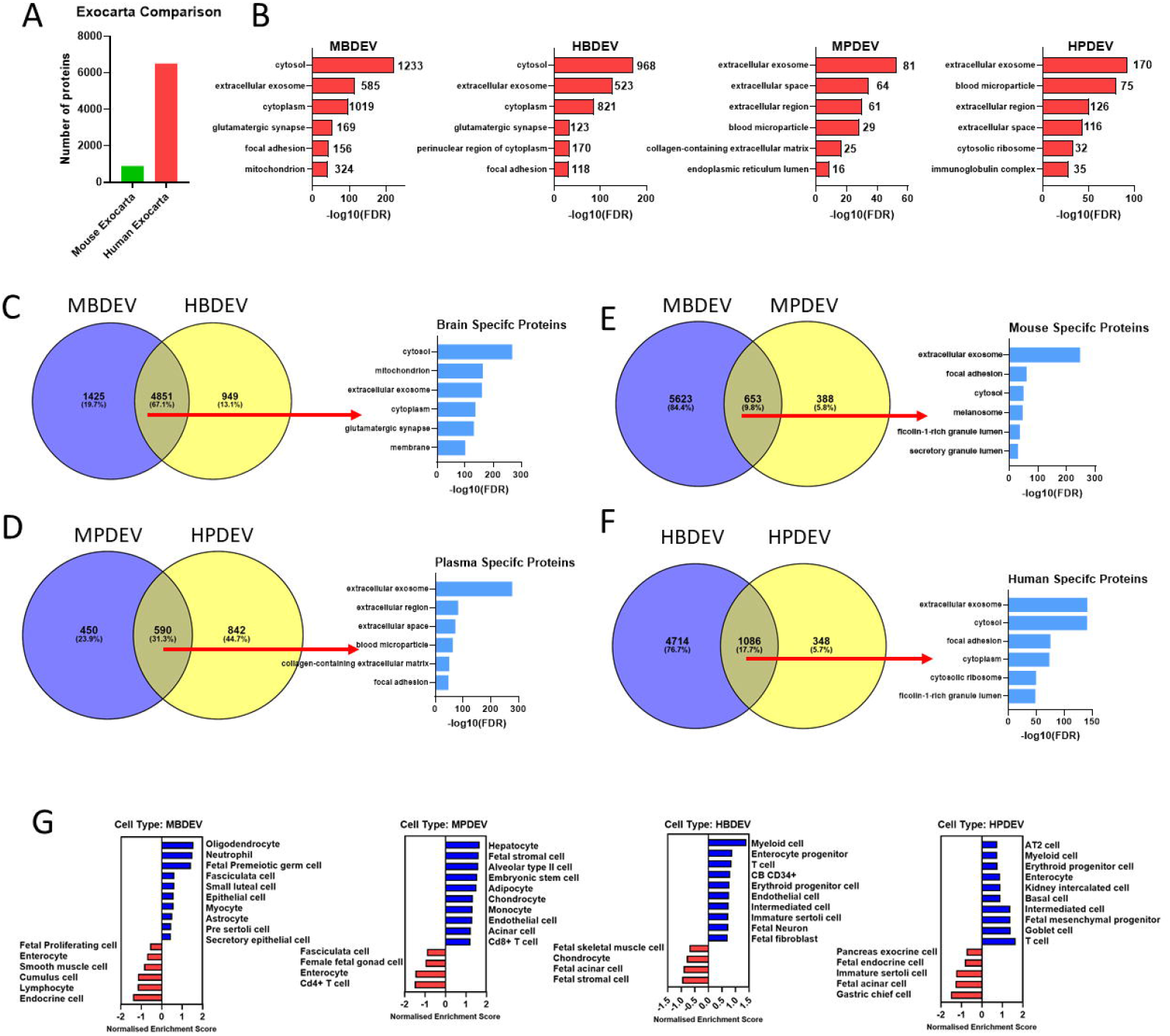
**(A)** Comparison of Exocarta protein annotations between mouse and human. **(B)** Top 6 GO: Cellular component terms associated with the reference sample protein signature from mouse brain, mouse plasma, human brain and human plasma. **(C-F)** Representation of overlapping proteins from MBDEV and HBDEV, MPDEV and HPDEV, MBDEV and MPDEV, and HBDEV and HPDEV related proteins and top 6 GO: Cellular component terms associated with the overlapping proteins. GO analysis was performed using DAVID. The bar plot displays the top six enriched GO terms, ranked by statistical significance. The y-axis represents the GO terms, while the x-axis shows the -log_10_(FDR), where higher values indicate stronger enrichment. **(G)** GSEA cell type enrichment analysis of MBDEV, MPDEV, HBDEV and HPDEV related proteins with WEB-based GEne SeT AnaLysis Toolkit. The bar plot displays the overrepresented (blue) and underrepresented (red) cell types, ranked by normalised enrichment score.

**Supplementary Figure 3.**
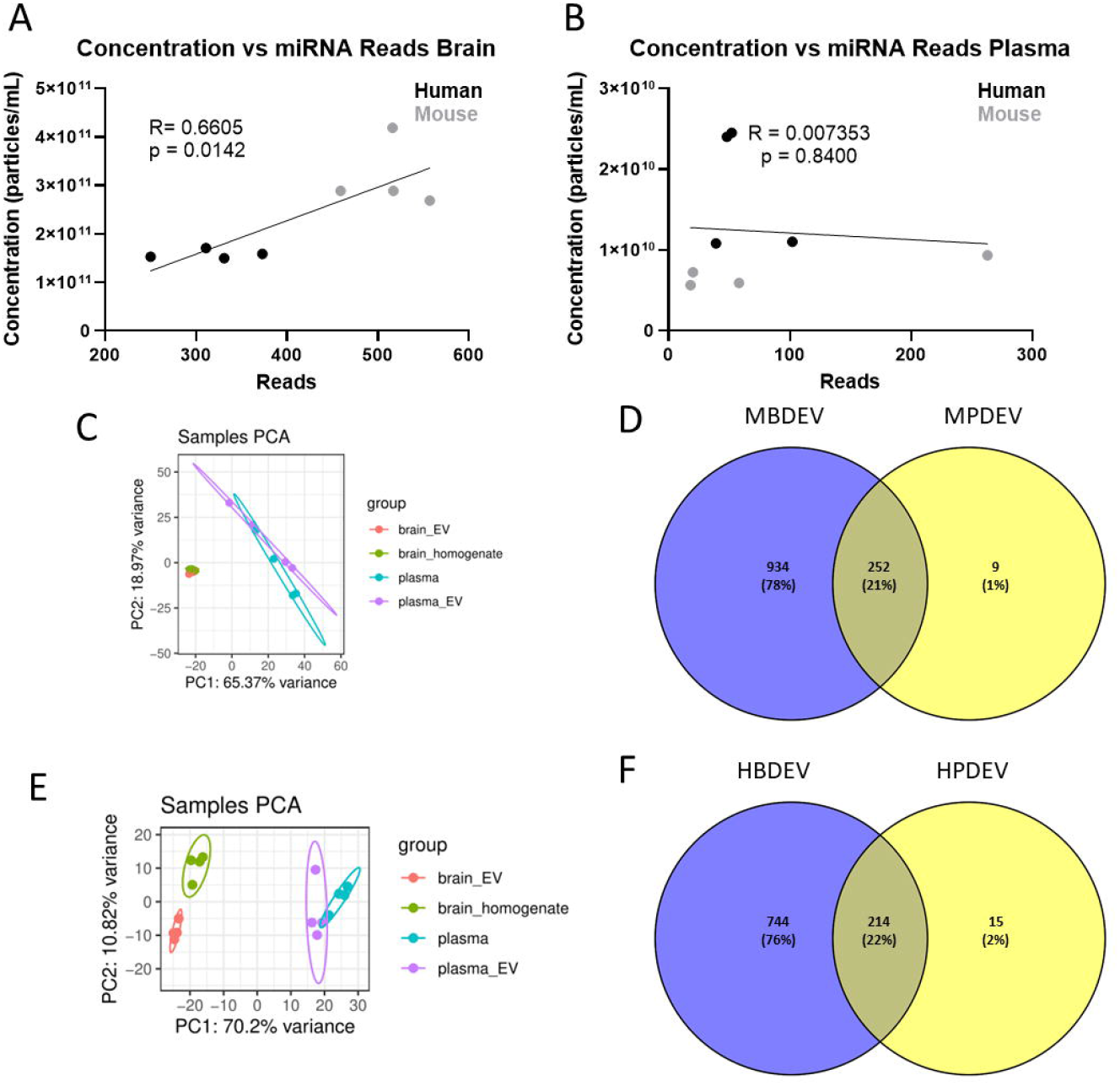
**(A-B)** Linear regression of the relationship between the starting concentration of EVs and resultant number of miRNA reads from brain and plasma. **(C)** PCA analysis of MBDEVs, MPDEVs, brain homogenate and plasma. **(D)** PCA analysis of HBDEVs, HPDEVs, brain homogenate and plasma. **(E-F)** Representation of overlapping miRNAs from MBDEVs and MPDEVs, and HBDEVs and HPDEVs.

**Supplementary Figure 4.**
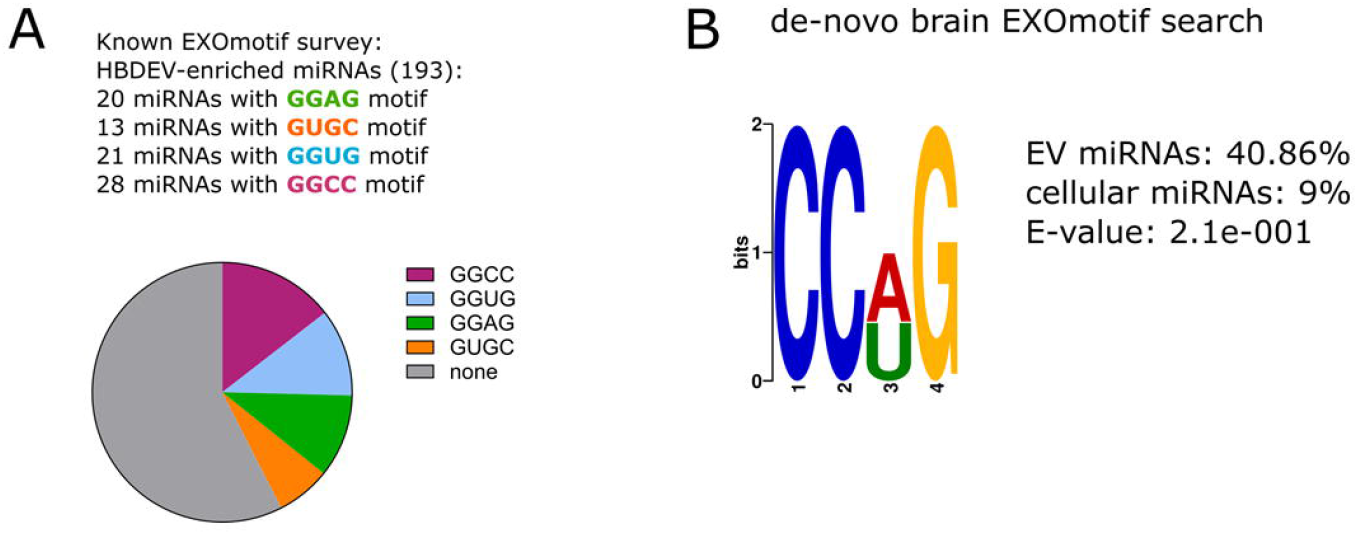
**(A)** Overview of known and identified EXOmotif survey of HBDEV-enriched miRNAs. **(B)** Search of de-novo EXOmotifs from HBDEVs (EV miRNA) and reference cell lysate (cellular miRNAs).

## Notes

### Competing Interest Statement

The authors have declared no competing interest.

